# Contradictory behavioral effects of neuronal perturbations on behavioral responses to linearly polarized light in freely walking *Drosophila*

**DOI:** 10.1101/2024.03.15.584848

**Authors:** Anna V. Titova, Andrew D. Straw

**Affiliations:** Institute of Biology I, Faculty of Biology, Albert-Ludwigs-Universität Freiburg; Freiburg, Germany; Bernstein Center Freiburg, Albert-Ludwigs-Universität Freiburg; Freiburg, Germany

## Abstract

Many insects can use the polarization of the skylight as a navigational cue. As shown previously, freely walking *Drosophila* orient along the e-vector of linearly polarized UV light presented both dorsally and ventrally. We are interested in the neuronal mechanisms leading to this behavior, and specifically how the central complex and its inputs are involved. We investigated the behavior of flies exposed to linearly polarized near-UV light (400 nm) presented dorsally. Flies walked freely in a circular, flat arena surrounded by a heat barrier. Using the GAL4-UAS genetic system, we drove the expression of the potassium inward rectifier KIR2.1 to perturb each of several different neuron types of the polarization vision pathway. Perturbing EPG compass neurons in the central complex slightly weakened average alignment and increased its variability. On the other hand, when two different GAL4 lines driving expression in the ER4m ring neurons, identified by connectomics as the major polarization inputs to the fly central complex, were perturbed, the alignment strength increased. A similar effect was observed when the inputs to ER4m, the TuBu_a_ neurons, were perturbed. We did not predict EPG and ER4m perturbations to cause opposite effects. Further investigation would be required to understand the physiological mechanisms of these contradictory behavioral effects.

## Introduction

Many insects use linear polarization of light as a navigational cue. The polarization pattern of the sky is one of the major compass cues alongside the Sun’s position and brightness gradient. Additionally, polarized light can be reflected off different surfaces, such as water, leaves, etc., and help the animals recognize such surfaces (Heinloth et al., 2018).

The fruit fly *Drosophila melanogaster* was shown to perceive both dorsally and ventrally presented polarized light (Stephens et al., 1953; Wernet et al., 2012; Wolf et al., 1980) and to use the skylight polarization to set their orientation (Weir and Dickinson, 2012). Walking flies tend to walk mainly along the E-vector of linear polarization, presented dorsally or ventrally (Wernet et al., 2012). Flying tethered flies, on the other hand, were shown to keep a constant arbitrary angle to the e-vector (Mathejczyk and Wernet, 2019; Warren et al., 2018). This ability to keep a constant arbitrary angle to a stimulus is reminiscent of menotaxis relative to intensity cues, which has been demonstrated in tethered setups on both flying and walking flies (Giraldo et al., 2018; Green et al., 2019; Haberkern et al., 2022). Here we ask: what is the role of “compass” EPG neurons and of their polarization-tuned inputs for freely walking flies to respond to linear polarization of dorsally presented near-UV light?

The angle of polarization (AoP) of dorsally presented light is detected by specialized photoreceptors in the dorsal rim area (DRA) of the eyes. The anterior visual pathway leads the information through the dorsal rim region of the medulla, anterior optic tubercle, and the bulb to the ellipsoid body in the central complex, giving input to the compass system (Hardcastle et al., 2021). Starting from the AOTU, two parallel pathways for polarized light signals were identified: through the anterior bulb and through the superior bulb. The TuBu_a_ (Tubercle->Bulb anterior) neurons relay the information from the anterior bulb to the ER4m ring neurons, which are thought to be the major polarization input to the EPG neurons in the ellipsoid body (Hulse et al., 2021). The superior bulb is thought to integrate both polarized and unpolarized signals, with some of the TuBu_s_ (Tubercle->Bulb superior) neurons showing polarization sensitivity with preferences for orthogonal AoPs in different hemispheres (Hardcastle et al., 2021). A subset of ER2 neurons, which receive inputs from TuBu_s_, was shown to have polarization sensitivity (Hardcastle et al., 2021). Here we explore the behavioral effects of neuronal perturbations within the two above-mentioned pathways.

First, we established an experimental assay to measure the behavioral responses to linearly polarized light presented dorsally in freely walking flies. As expected, wild-type flies walked on average in the direction of the E-vector, termed ‘alignment’. Next, we used genetic perturbation to alter the activity of several neural populations of the polarization pathway leading to the EPG neurons and looked for effects on the polarization behavioral responses. Using the Gal4-UAS expression system, we expressed inward rectifier potassium channel KIR2.1 in EPG, ER4m, ER2, and in two populations of TuBu neurons. This is expected to hyperpolarize, or “silence” the cells. None of those perturbations abolished alignment completely, but alignment strength was affected. In the case of disrupting the pathway through the superior bulb, we also observed a change in the preferred direction at the early exposure to polarized light.

## Results

### Walking flies align with linearly polarized light

First, we established an experimental behavioral paradigm and checked if the flies with unperturbed neural circuits behaved as expected from the literature (Wernet et al., 2012). Flies walked in a flat circular arena surrounded by a heat barrier, exposed to near-UV light (400 nm) passing a polarizer-diffuser pair of filters, which could be switched to make the light linearly polarized or unpolarized (Fig. 1A). We used a motor to set the angle of polarization. In every trial, for 20 minutes a group of flies was exposed to linearly polarized light with the direction changed by 90° every 5 minutes. Next, the light was turned off and the flies walked in darkness for another 5 minutes. As expected, we observed that the flies tend to align with the e-vector direction (Fig. 1B, C). When we repeated the same experiment with unpolarized light of the same intensity, by flipping the polarizer-diffuser pair of filters, the flies did not exhibit any strong directional preferences during the whole trial (Fig. 1D). In the polarized condition, the alignment strength increased over time: in the first stage the alignment was weak and saturated at a higher value in stages 3 and 4, corresponding to 10-20 minutes from the start of the experiment (Fig. 1F).

**Figure 1.**
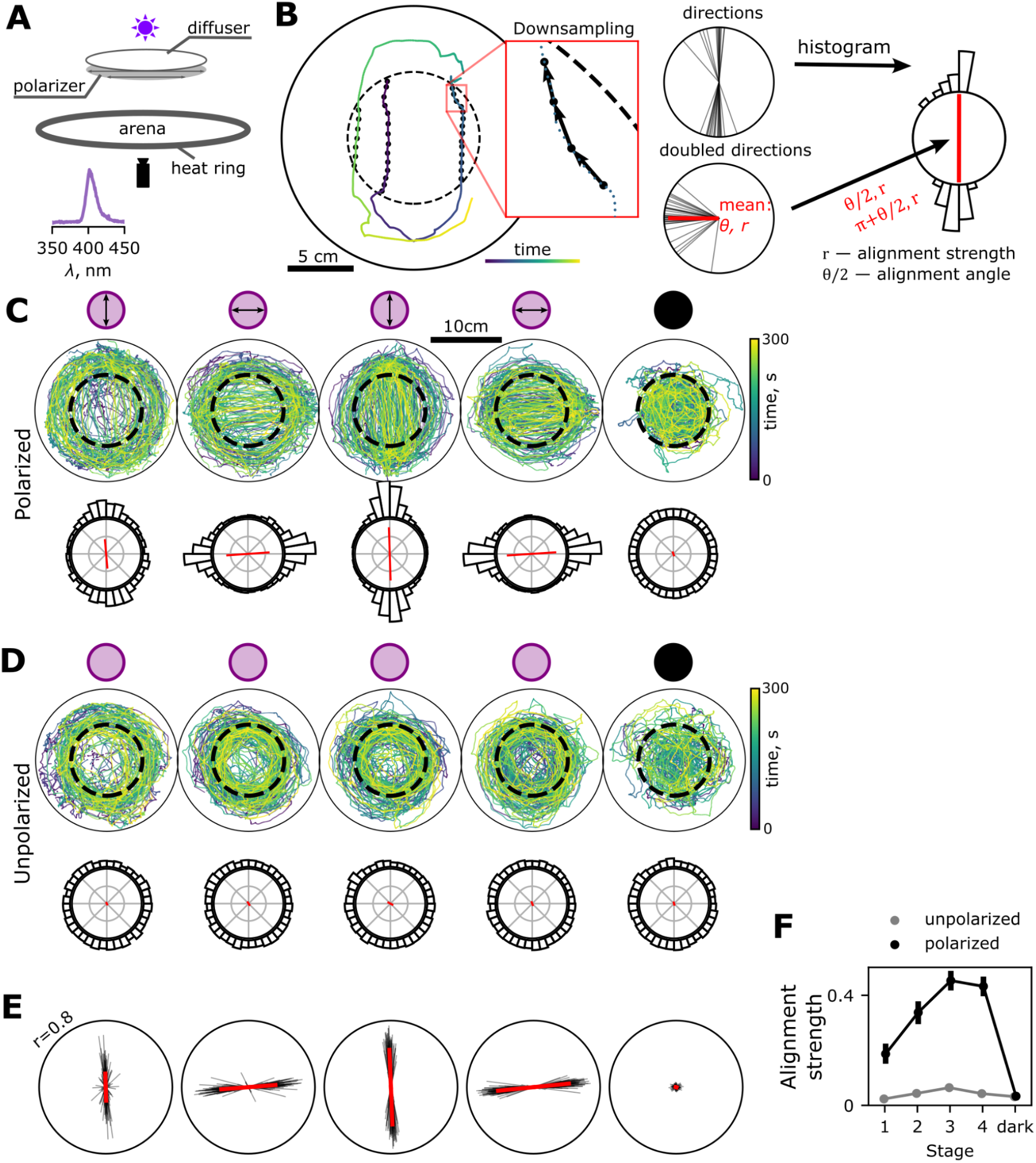
Walking flies align with polarized light. **A** Experimental setup. The flies are walking in a flat circular arena (20 cm diameter) with a heat barrier. Near UV light comes through a diffuser-polarizer pair that can be rotated by a motor. Bottom: LED spectrum measured at the arena center after passing the filters (peak intensity at 402 nm). **B** Processing illustration on a trajectory fragment shown on the left. Scale bar 5 cm. The trajectory is downsampled to 5 fps (black dots). For each point of the downsampled trajectory inside the central region (5 cm radius, dashed circle) the direction towards the next point is calculated. The distribution of directions is plotted as a circular histogram. The mean direction is calculated based on doubled values of directions and shown as a red line. Alignment strength is the length of the mean direction vector (from 0 to 1). **C, D** Examples of recorded group trajectories (5 flies in a group) in polarized (C) and unpolarized (D) conditions. Each stage is 5 minutes, the polarizer is rotated 90° between the stages. In the last stage, the light is off. The bottom row of each panel represents the distribution of walking directions in each stage acquired as shown in B. **E** Average directions of walking, each black line represents one group trial (4-6 flies), n=93 trials. The red line is the average vector of all trials. The circles correspond to an alignment strength value of 0.8. **F** Alignment strength stage by stage, measured as the vector strength of summed instantaneous directions over a stage (r value in B). Mean value between trials (n=93) and 95% bootstrapped confidence intervals for polarized condition (black), single values for unpolarized condition (gray).

### Perturbing neurons in the polarization pathway with Kir2.1 affects the polarization response

Next, we compared the behavior of different genotypes using this protocol. We genetically perturbed neurons along two polarization pathways (see Introduction) leading to the compass system. We used the GAL4-UAS system to express potassium inward rectifier Kir2.1 in populations of cells containing the neurons of interest (Fig. 2A) and compared the flies’ behavior between different genotypes (Figs. 2,3).

**Figure 2.**
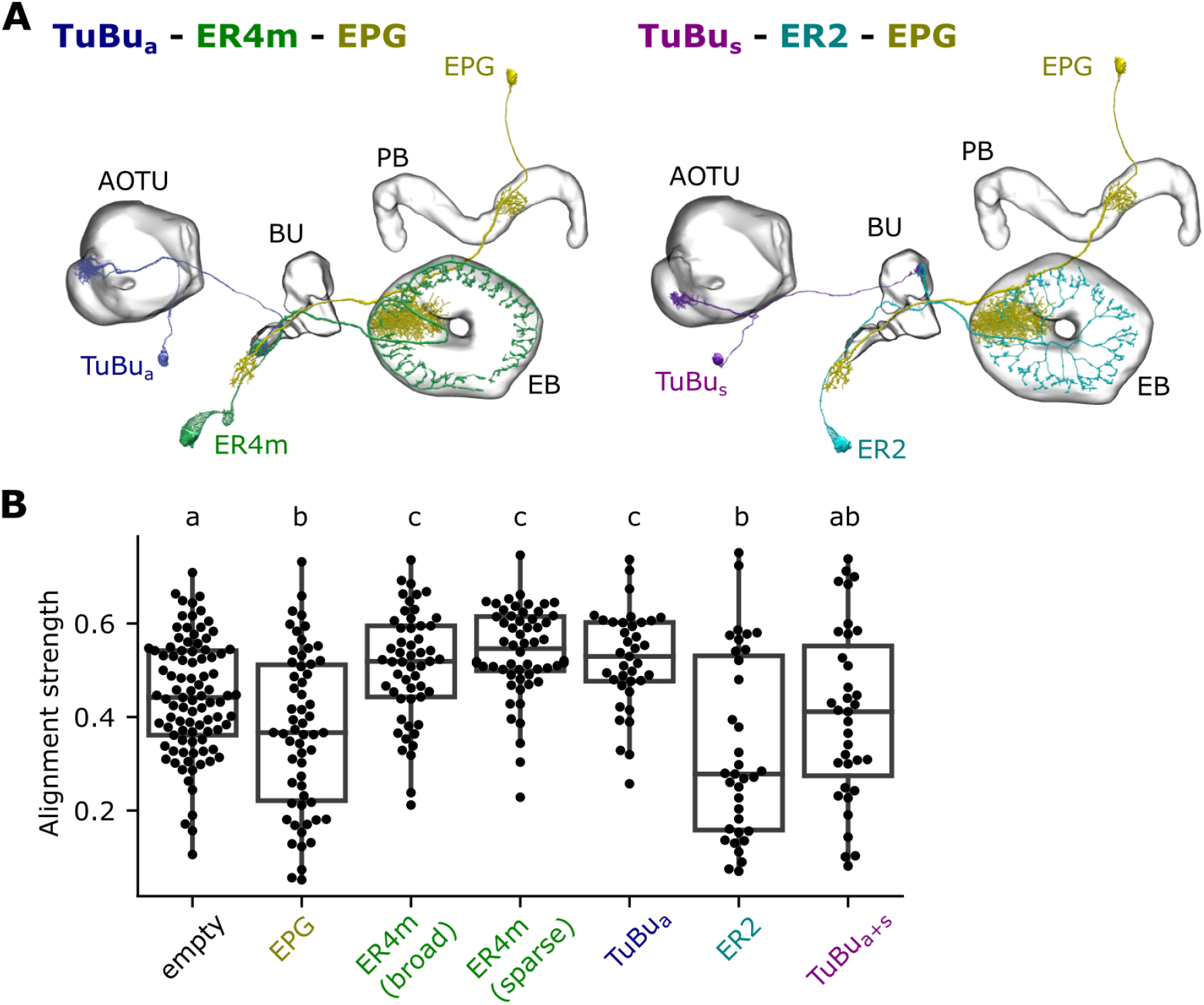
Effects of neuronal perturbation with Kir2.1 on the polarization response in walking flies. **A** Illustration of two UV polarization pathways to the compass (EPG) neurons, based on Hardcastle et al. (2021). Left: TuBu_a_ - ER4m - EPG; right: TuBu_s_ - ER2 - EPG. One neuron of each type is shown. Created using neuPrint web interface (Plaza et al., 2022). **B** Average alignment strength during the later half of 20-minute exposure to linearly polarized near-UV light by groups of 3-6 flies (N trials: empty: 93, EPG: 58, ER4m (broad): 53, ER4m (sparse): 56, ER2: 35, TuBu_a: 37, TuBu_a+s: 35). Alignment strength is measured as the length of the average instantaneous moving direction vector in the central part of the arena (see Fig.1B). Each dot represents one group trial. The trials are grouped by genotype, the labels correspond to the Gal4 line driving expression of Kir2.1. Different letters (a, b, c) indicate non-intersecting 94% HDI of posterior estimates of the means (see Methods and Fig.S2-1 for the details). See also Fig. S2-3 for alignment strengths dynamics from stage to stage.

**Figure 3.**
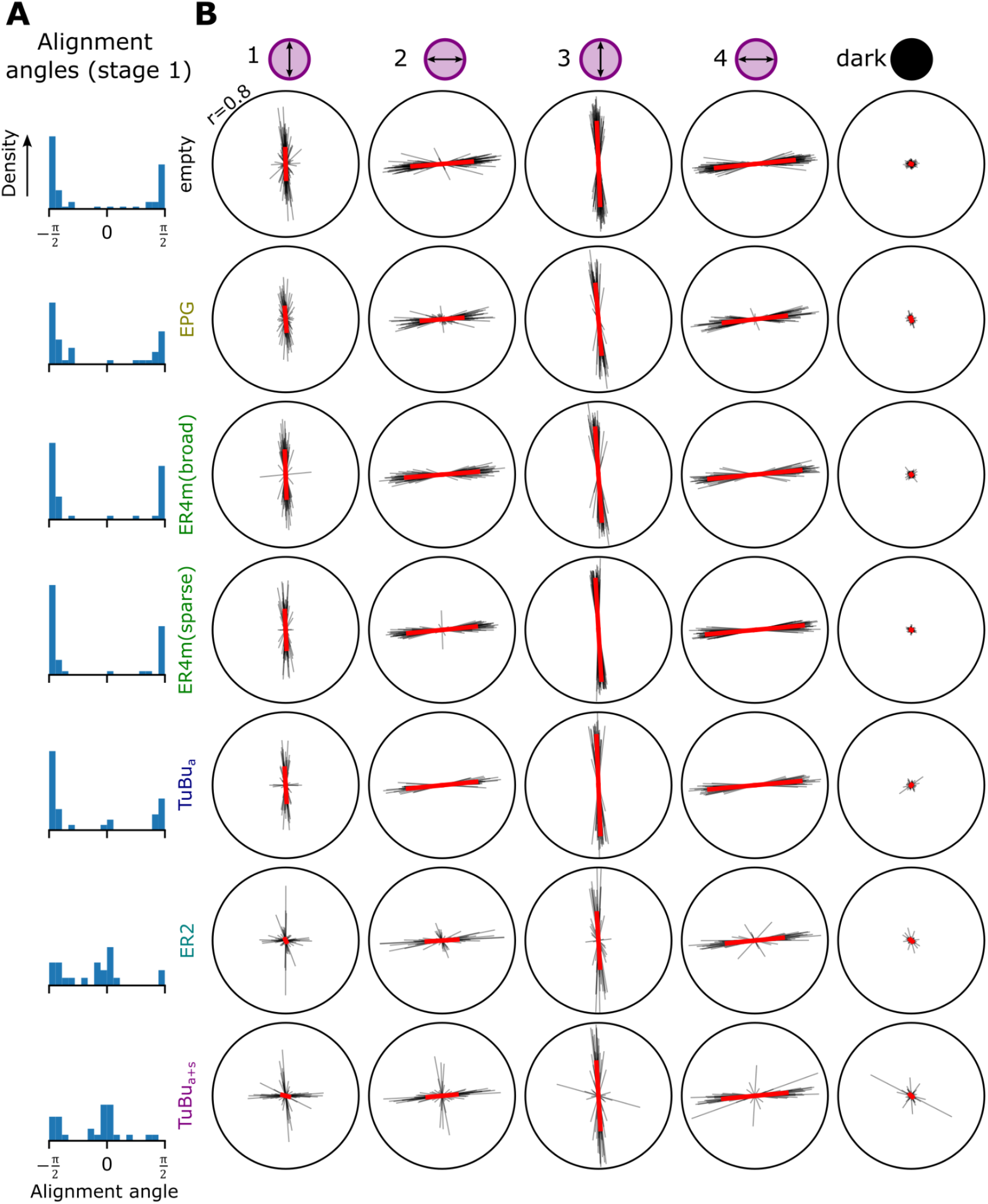
Perturbation of the ER2 pathway with Kir2.1 leads to bimodal heading preference at early exposure to linearly polarized light. **A** Distributions of alignment angles in the first stage (5 minutes, polarizer orientation π/2). The calculation of alignment angles was based on doubled moving direction values, so the range is half the circle. Only vectors with lengths above the threshold corresponding to 95% quantile of alignment strengths in the dark stage contribute to the distributions. In ER2 and TuBu_a+s_, around half of the alignment angles are close to 0 (perpendicular to the polarizer, Fig.S3). **B** Average walking directions. Each row represents data for perturbing one neuron population, columns corresponding to 5-minute stages of the experiment with polarization direction changed 90° between stages. Each black line represents the average direction of walking for one group trial (3-6 flies, see Fig.1B and Methods). The red line is the vector average of the group trials. Due to polarization having a periodicity of π radians (180°), vectors are plotted at their original angle and with an additional angular offset of π.

When we perturbed EPG neurons (SS00096>Kir2.1), we observed a slight decrease in alignment strength, coupled with increased variability (Fig.2B, Fig S2-1). Assuming a reliable inhibition of EPG neurons, this suggests that an EPG-independent pathway is largely sufficient for alignment, but that the EPG pathway also contributes to alignment.

Previous studies reported that the polarization inputs to the central complex come from two populations of ring neurons: ER4m is the major polarization input according to connectomic work (Hulse et al. 2021), and some ER2 neurons also show polarization tuning (Hardcastle et al., 2021). We targeted these two ring neuron types and corresponding subtypes of TuBu neurons projecting on them. (To target ER2 neurons, we used R18C09-GAL4. Hardcastle et al., 2021 attribute physiologically measured polarization responses with this line to the ER2 neurons but note that this line also drives expression in ER3w and ER3p. Garner et al. 2023 report no polarization inputs to ER2 based on a connectomic analysis but indicate ER3w receives polarization input. We follow Hardcastle’s nomenclature and use the name ER2.)

We observed an increase of alignment when the ER4m neurons were perturbed, using two different Gal4 lines. A similar effect was caused by perturbing TuBu_a_ neurons, which were shown to convey polarization input to ER4m neurons (Hardcastle et al., 2021). The similarity between TuBu_a_ and ER4m perturbation was expected and supports the hypothesis that the effect is related to the polarization information carried by both neuron types. Increased polarization alignment was, however, unexpected given our findings with EPG perturbation. We predicted that if the EPG neurons positively contribute to alignment, then perturbing the major polarization input to them would reduce their polarization-tuned contribution and therefore the alignment strength.

Perturbation of ER2 neurons led to a slight decrease of the average alignment strength, similar to the EPG perturbation. To perturb TuBu_s_ neurons, we used the driver line that targets TuBu neurons with processes in both the anterior and the superior bulb. Their perturbation did not lead to significant alignment strength changes. Additionally, we observed an interesting effect on directions of alignment: in the earlier trial stages (stage 1 and sometimes stage 2), flies with perturbed activity of ER2 or TuBu_s,a_ neurons sometimes aligned with a direction perpendicular to the e-vector of polarization. In later stages, however, this bimodality of alignment angles disappeared (Fig. 3, Fig. S3).

Further, we compared the variances of alignment strength values for the different genotypes. Alignment strength in late stages (stages 3-4 of polarized light exposure) was more variable for EPG>Kir2.1 and ER2>Kir2.1 than for controls, ER4m>Kir2.1, and TuBu_a_>Kir2.1 flies (Fig. S2-1). The effects of EPG and ER2 perturbation on the alignment strength are statistically similar.

## Discussion

In this study, we established an experimental paradigm that allows the characterization of the behavioral response to linearly polarized light presented dorsally in walking flies by analyzing their trajectories. As expected based on previous work, the flies aligned with the e-vector of linear polarization in the control condition, and alignment strength increased over time. Perturbing EPG compass neurons with genetic expression of the potassium inward rectifier Kir2.1 led to a slight decrease in alignment strength and an increase in variability between trials. Perturbing the ER4m pathway with Kir2.1 led to a surprising increase in alignment strength. Perturbing the ER2 pathway changed alignment angle in early, but not late, stages of polarized light exposure.

It may be informative to compare fly behavioral responses to polarized light with the well-studied behavioral responses of flies to luminance-defined stripes. Neurons involved in the behavioral responses to both types of stimuli may be shared, such as in areas downstream from raw visual processing like the central complex. Tethered flies can “fixate” stripes at angles near zero or they can exhibit “menotaxis” during which the stripe is held at near constant, but non-zero angle (Heisenberg & Wolf, 1984, Giraldo et al., 2018, Green et al., 2019). Fixation may indicate an attempt to approach the object, whereas menotaxis would be expected if the fly was attempting to travel at a fixed compass direction relative to a distant landmark for a sustained period. Menotaxis-like responses to linearly polarized light have been demonstrated in some of the previous studies in tethered flying flies (Warren et al. 2018, Mathejczyk and Wernet, 2019). The polarization alignment behavior measured in this study and prior studies (e.g. Wolf et al, 1980, Wernet et al. 2012) shows interesting similarities to stripe fixation, as flies locomote with an angle of near zero relative to the e-vector of polarized light. The causes underlying fixation-like or menotaxis-like responses are not known to us, and some changes in our experimental protocol could potentially lead to menotaxis-like responses to linear polarization in walking flies.

Previous studies showed that EPG neurons are required to perform menotaxis in tethered setups and that their silencing leads to fixation behavior (Giraldo et al., 2018; Green et al., 2019; Haberkern et al., 2022). Here, when we perturbed EPG neurons (albeit from a condition that more resembles stripe fixation than menotaxis as described above), we observed a slight decrease in alignment strength and an increase in variability between trials. The simplest explanation is that the polarization alignment we recorded is not akin to menotaxis and that EPG perturbation reduces a fly’s ability to hold a straight course, even a zero-angle course relative to the e-vector of polarized light. Given that EPG perturbation only resulted in a partial reduction of polarization alignment, two non-exclusive hypotheses arise. First, our genetic perturbation technique may have been only partially effective at interfering with EPG activity. Second, an EPG-independent pathway may play a substantial role in mediating polarization alignment responses.

The increase in alignment following the genetic perturbation of ER4m inputs to the CX was unexpected, given that EPG perturbation did not lead to the same outcome. We expected that the effects of perturbing EPG and ER4m neurons would be similar, given that ER4m neurons provide the strongest polarization input to EPGs(Hardcastle et al., 2021; Hulse et al., 2021). Hypothetically, silencing of the inhibitory ER4m neurons could lead to disinhibition of other ring neurons that receive inputs from ER4m, thereby increasing their influence on the EPG activity. Given that there are other polarization-sensitive ring neurons (a subset of ER2 or ER3w) that receive inputs from ER4m neurons (Garner et al., 2023; Hulse et al., 2021), their signal could get enhanced and the resulting behavioral response could become stronger. Additional experiments measuring neural activity are necessary to test this physiologically.

## Methods

### Fly rearing and genotypes

We used 5-7 days-old mated female flies for the experiments. The flies were reared at 25°C, 60% humidity with a 12h:12h light:dark cycle with lights on at 8 am. A day before the experiment the flies were briefly anesthetized at 4°C, sorted by gender, and the female flies were starved for 20-25 hours in a vial with water-soaked paper.

To obtain the experimental genotypes we crossed Gal4 lines from Table 1 with w^+^; UAS-mKir2.1-T2A-tdTomato; tub-GAL80^ts^ to express the modified mouse Kir2.1 in the cells of interest (Vijayan et al., 2023). The UAS line was obtained from Gaby Maimon and corresponds to the Bloomington stock number 600374. We used the same GAL4 lines that were used in the polarization pathway study of Hardcastle et al. (2021). Below we list correspondences between GAL4 lines and the neuron types in the hemibrain connectome (Scheffer et al., 2020), as reported in Hardcastle et al. (2021). The TuBu_a+s line (R88A06) contains TuBu01, TuBu06, TuBu08 and TuBu09 neuron subtypes. According to Hardcastle et al. (2021), the ER2 line R19C08 contains four subtypes of ER2 neurons (ER2_a, ER2_b, ER2_c, ER2_d) and additionally ER3w neurons and ER3p neurons. They reported ER2_a, ER2_b, and ER2_d as potentially polarization-sensitive. However, as suggested later (Garner et al., 2023), it could be that ER3w neurons, also labeled by R19C08, are polarization sensitive and potentially encode both polarization and chromatic information of the sky. The TuBu_a_ line R34H10, among others, contains TuBu01 neurons, which form 1-to-1 connections to ER4m neurons in the anterior bulb.

**Table 1:** GAL4 and split-GAL4 lines used in the study. To obtain the experimental genotypes, virgin females from the GAL4 lines were crossed to males of genotype w^+^; UAS-mKir2.1-T2A-tdTomato; tub-GAL80^ts^ (BDSC-600374). In the text, the experimental genotypes are referred to as X>Kir2.1, where X is the short name from this table. * According to Hardcastle et al. (2021), GMR19C08-GAL4 also drives expression in ER3w and ER3p neurons.

| Short name | Stock number (Bloomington) | Genotype |
| --- | --- | --- |
| empty | BDSC-68384 | w[1118];;GAL4.1Uw |
| EPG |  | w;19G02-p65ADZp; 70G12-ZpGdbd (SS00096) |
| ER4m (sparse) | BDSC-39209 | w[1118];;GMR59B10-GAL4 |
| ER4m (broad) | BDSC-49784 | w[1118];;GMR34D03-GAL4 |
| ER2* | BDSC-48845 | w[1118];;GMR19C08-GAL4 |
| TuBu<sub>a</sub> | BDSC-49808 | w[1118];;GMR34H10-GAL4 |
| TuBu<sub>s+a</sub> | BDSC-46847 | w[1118];; GMR88A06-GAL4 |

We verified the expression by cutting a rectangular window in the head and checking fluorescence (data not shown) in 2-3 flies of each experimental genotype. The presence of the GAL80^ts^ construct could lead to lower expression of Kir2.1 in the targeted cells, when the flies were raised at 25°C, which is less than the permissive temperature stated for GAL80^ts^. We could observe the fluorescent cells in the experimental flies, which confirmed the presence of Kir2.1. Without the GAL80^ts^ construct crosses with two of the GAL4 lines produced mostly lethal phenotypes (with ER4m sparse and TuBu_a_ lines). Moreover, we observed a difference in the empty-Gal4 controls behavior: the average alignment was weaker for the empty>Kir2.1 than for empty>Kir2.1, Gal80^ts^ flies (data not shown), which might be explained by a leaky expression of Kir without the Gal80^ts^ suppressor. Taking this into account, we decided to proceed with using the UAS-Kir2.1, Gal80^ts^ line and raising the flies at 25°C.

The experiments were conducted in the late afternoon (15:00-20:00h). The temperature in the experimental room was maintained at 23°C, but the temperature inside the arena was higher because of the heat barrier. To transfer the flies from the vial to the arena, we applied airflow through a tube attached to a pipette tip with a cut end. The recording started soon after the flies were transferred to the arena (around a 1-2-minute delay due to loading the flies to two setups in parallel and adjusting tracking before starting a recording). Between the trials, the arena floor was wiped with 75% ethanol.

### Experimental setups

We used two similar setups in parallel for the experiments. The 20 cm walking arena, illuminated by an infrared LED strip located on the sides (Fig. 1A, Fig. S1A) was placed on a lifted transparent platform inside a light-tight box. The tracking was performed using an infrared-sensitive camera located under the arena and facing up. Between the arena and the transparent plate, we put a dark red filter to minimize the amount of reflections the flies can see from below. Above the arena, we placed a custom-made polarized light illumination mechanism with a stepper motor and a gear system to rotate the polarizer (Fig. S1D). In one setup we used the linear polarization filter PL 55 HOYA. We used a linearly polarized film filter for the second setup because the same model was out of stock, leading to differences in light intensity between setups (Table 2, Fig. S1B,C). We used the projection screen material (Gerriets International Opera Creamy White Front/Back) as a diffuser in both setups.

**Table 2.** Polarized light intensity, measured in two experimental setups. Intensity in the right column was measured in the arena center. See also Fig. S1.

| Setup | Lambda_max, nm | FWHM, nm | Intensity, $\mu\text{W}/\text{cm}^2$ |
| --- | --- | --- | --- |
| Setup #1 | 404.4 | 15.25 | 0.966 |
| Setup #2 | 401.8 | 15.62 | 1.453 |

The intensity at 400 nm was measured using the power meter PM100A (Thorlabs) and the laser power sensor S130C (Thorlabs). The spectra were measured in arbitrary units using the Ocean Optics spectrometer (model: USB2000+VIS-NIR).

### Experimental procedure and tracking

Each trial was performed on a group of 4-6 flies of the same genotype. Before the recording, the UV LED was off. The flies were transferred to the arena under dim light, the setup lid was closed and the recording was started. A trial consisted of 5 stages, 5 minutes each.

During the first 4 stages, the UV LED was on, between the stages the filter was rotated by 90°, therefore changing the orientation of the e-vector. The direction sequence was similar in all trials. During the last 5-minute stage the light was off.

The tracking was performed by Strand Camera software (https://strawlab.org/strand-cam) at 30 Hz. We used a Basler Ace acA1300-200um camera with a Tamron 12VM412ASIR lens and an infrared filter (Lee Infrared Transmission 87 Filter), providing infrared illumination from the sides with an IR LED strip. The tracked recording consists of 2D coordinates of detected objects at each frame and their estimated velocities. The individual trajectory segments are marked as the same object, but the collisions between objects are not resolved, so we cannot reliably extract the whole trajectory of an individual fly.

### Data analysis

#### Alignment strength and angle

The calculation of alignment strength and angle is illustrated in Figure 1B.

The tracked recordings (description in Methods: Experimental procedure and tracking) were downsampled from 30 Hz to 5 Hz. The trajectory segments shorter than 30 detections (1 s) were removed before downsampling. Downsampling was performed for each trajectory segment by taking median x and y coordinates in each time step of 0.2 s.

Then speed and direction of movement were calculated based on downsampled data using the coordinates of two adjacent points. The points with a speed less than 0.005 m/s were labeled as stops and excluded from further analysis.

The alignment was calculated in the central region of the arena — a circle with a 5 cm radius — ignoring the outer region, where the flies perform turns due to heat at the border. Because polarized light has a period of pi radians (180°), we doubled the angular directions of movement at each point and viewed these as unit vectors (with length one and the corresponding direction). We then calculated the alignment vector as a vector average of doubled directions (with direction θand length *r*). We define the alignment strength as the length of this vector, *r*, with a value between 0 and 1, where 0 means no preference for any angle and 1 means always following the same direction or its direct opposite. The alignment angle is defined as θ/2. A certain alignment angle means that the flies walk preferably in this direction or the opposite direction (π + θ/2). Note that alignment strength does not include any knowledge about direction and only characterizes the amount of movement in a preferred direction.

#### Comparison of alignment strengths distributions

We applied Bayesian model fitting using Markov-Chain Monte Carlo software (PyMC 5.1.1) to compare the distributions of alignment strengths between different genotypes. We model the alignment strengths for each genotype as a beta distribution with parameters α_*x*_ and β_*x*_, where X corresponds to the driver line (genotype X-GAL4>UAS-Kir). We used identical, uninformative priors for each genotype:

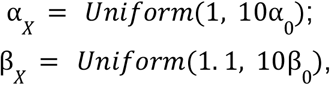

where α_0_ and β_0_ were calculated based on the distribution of vector strengths across all genotypes pooled

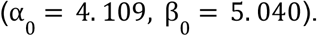

After fitting the models with observed data, we compared the 94% highest density intervals (HDI) of posterior estimates of the means and variances between different genotypes (Fig. S2-1). The letters on top of Fig. 2B indicate non-intersecting HDIs, corresponding to a reliable difference between vector strengths of genotypes assigned different letters. The resulting HDI intervals for means and variances of alignment strengths are presented in Supplementary Table 1.

#### Alignment angles comparison

We classified the alignment angles in the first stage into two categories: around 0 and around π/2 (90°) – thus the doubled angle falls into categories around 0 and around π: − π/2 < θ< π/2 and π/2 < θ< 3π/2. We modeled the alignment angles as an independent Bernoulli distribution for each genotype. In this model, the Bernoulli distribution parameter *p* is the probability that the alignment angle is in the first category (around 0). If *p* is close to 0, it means that most alignment angles of the first stage are in the second category (around π/2), and thus close to the e-vector. A value of *p* 0.5 means that in half of the trials, the flies aligned in the direction perpendicular to the e-vector. A value of *p* 1 means that all flies were of the first category, and thus aligned perpendicular to the e-vector. The priors for *p* were uniform distributions from 0 to 1 and were identical for each genotype. After fitting the models with observed data, we compared 94% higher density intervals (HDI) of posterior estimates of *p* between different genotypes (Fig. S3).

## Supplemental info and figures

**Figure S1.**
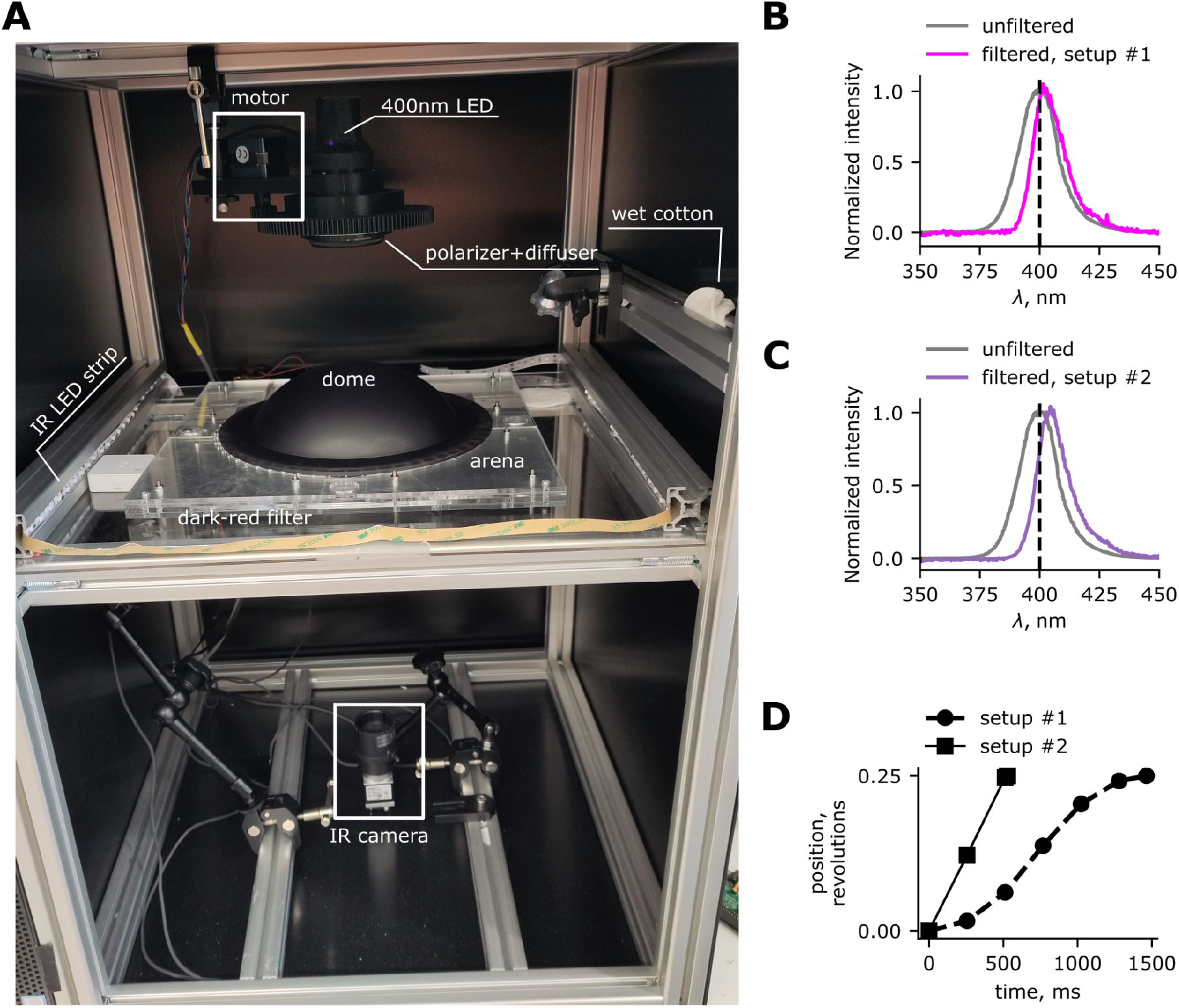
Experimental setups. **A** Photo of the experimental setup with labeled components. **B, C** Normalised spectra of the LEDs measured without filters and with filters. The unfiltered LED spectrum shown on both panels was measured in setup #2. **D** Motor rotation profile of both setups (different due to different motor models used), 90° (0.25 revolutions) rotation between the experiment stages.

**Figure S2-1.**
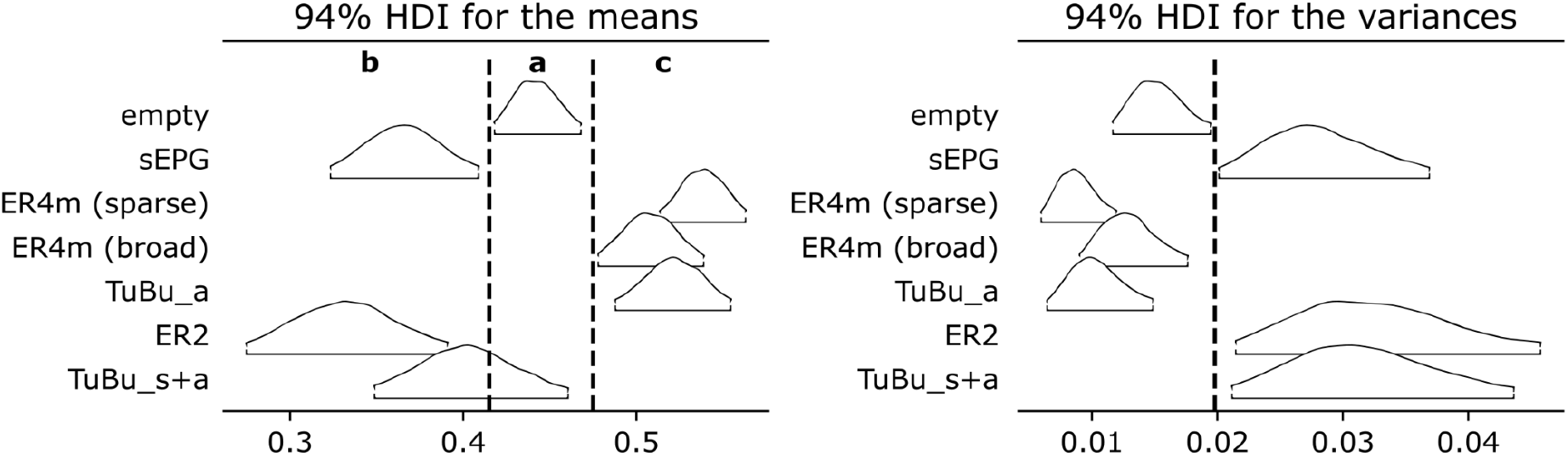
94% HDI of posterior estimates of the means and variances of alignment strengths. The vertical lines indicate the borders of non-intersecting interval groups, labeled a,b,c, as in Fig.2.

**Figure S2-2.**
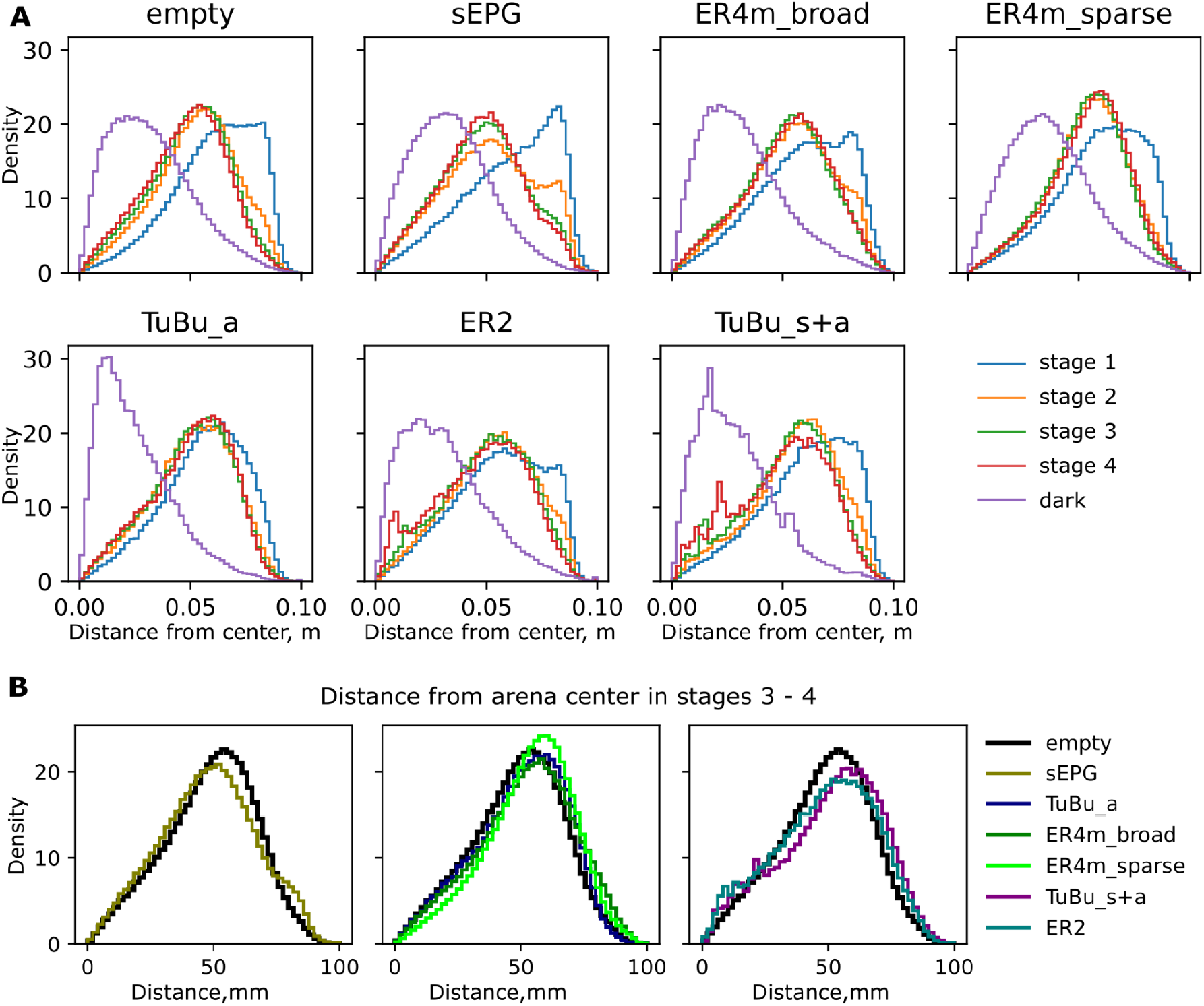
Distributions of distances from the arena center. **A** Each panel represents the average distributions of one genotype (X>Kir2.1, where X is the panel tile), series represent different stages of the experiment: stages 1-4: polarized light is on, dark: light off. **B** Average distributions of distances in stages 3 and 4 combined. Left: empty>Kir2.1, sEPG>Kir2.1; center: empty>Kir2.1, TuBu_a>Kir2.1, ER4m>Kir2.1, right: empty>Kir2.1, TuBu_s+a>Kir2.1, ER2>Kir2.1. Alignment strength in Fig.2 was calculated based on the trajectories in stages 3 and 4, where the distance to the center was less than 50 mm.

**Figure S2-3.**
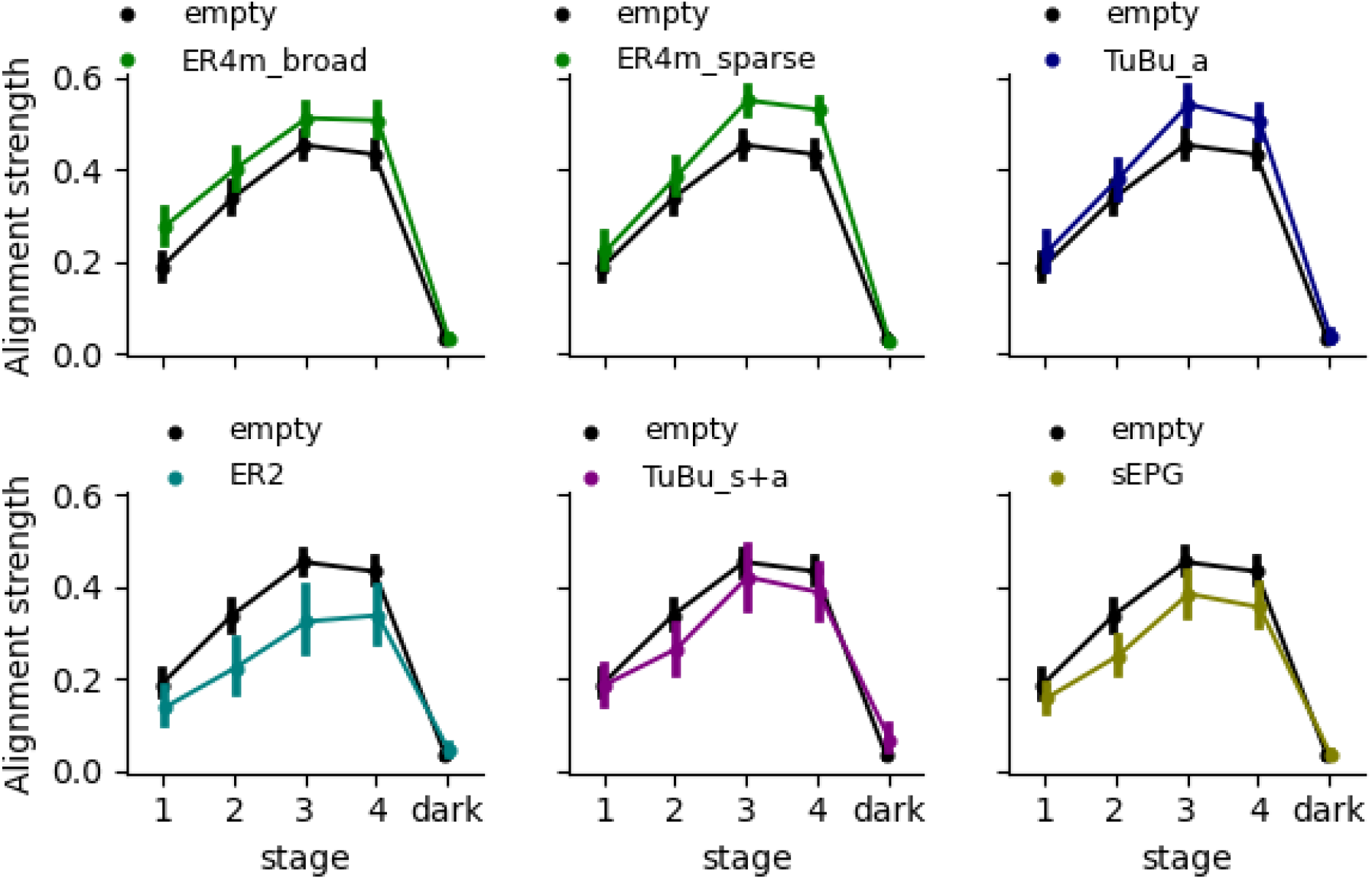
Stage-by-stage alignment strength of flies with perturbed polarization vision pathway. Genotypes are X-Gal4, UAS-Kir2.1, where the driver X is specified in the legend (see Table 2 for exact driver lines).

**Figure S3.**
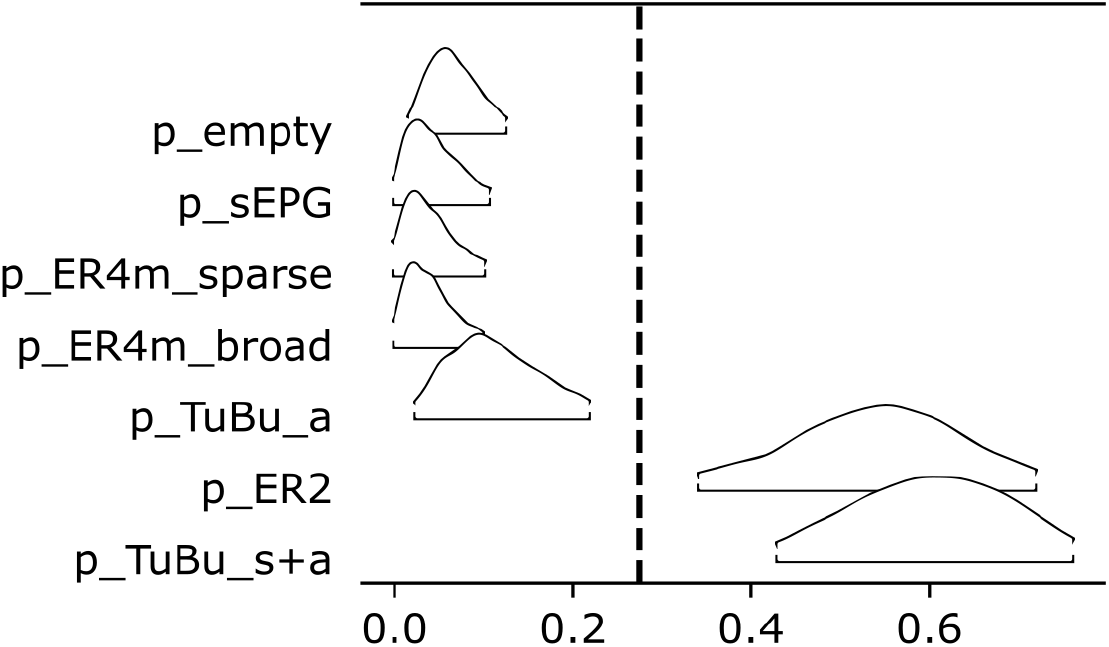
HDI of posterior estimates of the probability of alignment to the direction perpendicular to the polarizer in the first stage of the experiment. The alignment angles for genotype X-Gal4>UAS-Kir2.1 are modeled as Bernoulli trials with probability p_X indicating the probability that the alignment angles were near 0 (meaning, the flies aligned perpendicular to the e-vector). Only trials with alignment strength above 95% quantile of the dark stage contribute to the analysis (same as in Fig.3A). The vertical line indicates the border of non-intersecting interval groups.

**Supplementary Table 1.** 94% HDI of the means and variances of alignment strengths in the later stages of polarized light exposure.

| HDI of posterior estimates of the means |  |  | HDI of posterior estimates of the variances |  |  |
| --- | --- | --- | --- | --- | --- |
| X-GAL4>UAS-Kir2.1 | hdi_3% | hdi_97% | X-GAL4>UAS-Kir2.1 | hdi_3% | hdi_97% |
| empty | 0.41668 | 0.46542 | empty | 0.01149 | 0.01899 |
| EPG | 0.32373 | 0.40618 | EPG | 0.02077 | 0.03702 |
| ER4m (sparse) | 0.51406 | 0.56221 | ER4m (sparse) | 0.00618 | 0.01177 |
| ER4m (broad) | 0.47923 | 0.53819 | ER4m (broad) | 0.00891 | 0.0175 |
| TuBu_a | 0.48831 | 0.55242 | TuBu_a | 0.00655 | 0.01479 |
| ER2 | 0.2739 | 0.38871 | ER2 | 0.02163 | 0.04527 |
| TuBu_s+a | 0.34825 | 0.45968 | TuBu_s+a | 0.02085 | 0.04315 |

## Acknowledgments

We thank T. Thang Vo-Doan for his help with building the experimental setup, Viktor Titov for helping with the spectral measurements, Mathias Wernet for valuable discussions concerning this study, and Gaby Maimon for sharing the UAS-Kir2.1 line we used. We thank Bloomington Drosophila Stock Center for providing other transgenic fly lines used here.

This work was funded by DFG grant STR 1357/6-1 to ADS.

## Author contributions

AVT and ADS conceived the study. ADS developed the tracking software, AVT developed software to run the experiment. AVT assembled the experimental setup and performed the experiments. AVT analyzed the data with input from ADS, ADS performed the statistical analysis of alignment angles. AVT and ADS wrote the manuscript.

## Data availability

Data and code are available upon request to the authors. The code used to perform statistical comparisons will be publicly available at the time of publication.

